# The Starlet Sea Anemone, *Nematostella vectensis*, possesses body region-specific bacterial associations with spirochetes dominating the capitulum

**DOI:** 10.1101/2020.05.10.084863

**Authors:** A. M. Bonacolta, M. T. Connelly, S. Rosales, J. del Campo, N. Traylor-Knowles

## Abstract

Sampling of different body regions can reveal highly specialized bacterial associations within the holobiont and facilitate identification of core microbial symbionts that would otherwise be overlooked by bulk sampling methods. Here we characterized compartment-specific associations present within the model cnidarian *Nematostella vectensis* by dividing its morphology into three distinct body regions. This sampling design allowed us to uncover a capitulum-specific dominance of spirochetes within *N. vectensis*. Bacteria from the family Spirochaetaceae made up 66% of the community in the capitulum, while only representing 1.2% and 0.1% of the communities in the mesenteries and physa, respectively. A phylogenetic analysis of the predominant spirochete sequence recovered from *N. vectensis* showed a close relation to spirochetes previously recovered from wild *N. vectensis*. These sequences clustered closer to the recently described genus *Oceanispirochaeta*, rather than *Spirochaeta perfilievii*, supporting them as members of this clade. This suggests a consistent and potentially important association between *N. vectensis* and spirochetes from the order Spirochaetales.

## Introduction

The discovery of microniche specific microbial communities has become important for our understanding of microbial interactions and diversity (Ainsworth, Thurber and Gates 2010; Bourne, Morrow and Webster 2016). Microbial niches have been identified in many different organisms from an ecosystem scale, all the way down to the cellular level (Aagaard *et al*. 2014; Foster *et al*. 2017; Ricci *et al*. 2019). For example, in the Giant Clam, *Tridacna maxima*, different tissues harbor distinct microbial communities, with the gills harboring a high prevalence of *Endozoicomonas* bacteria. Compartmental evaluation of *T. maxima* helped uncover *Endozoicomonas*’s role in nitrogen-cycling within the holobiont (Rossbach 2019). The microniches within animal tissues are proposed to strongly influence the composition of the microbial community (Vaishnava *et al*. 2008; Hooper, Littman and Macpherson 2012). Consequently, the specific composition of each microniche can provide clues as to the community’s functional contribution to host-physiology (Ainsworth, Thurber and Gates 2010).

In the Phylum Cnidaria (sea anemones, corals etc.) many species are known to associate with different symbiotic microbial partners, including bacteria, and these relationships are known to be important for the health of the cnidarian host (Rädecker *et al*. 2015; Bourne, Morrow and Webster 2016; Robbins *et al*. 2019). Previous studies have identified that whole organism sampling methods used for microbiome analysis can skew results by preferentially sampling certain compartments or by combining them and drawing conclusions based on the agglomeration (Sweet, Croquer and Bythell 2011). An advantage of compartmental sampling of an organism’s body is that it can facilitate the identification of core microbial symbionts in different body compartments, which likely represent highly specialized symbiotic associations (Apprill, Weber and Santoro 2016; Bourne, Morrow and Webster 2016). However, in the model sea anemone, *Nematostella vectensis*, the presence of body region-specific associations of microbial communities has not previously been identified.

*N. vectensis* is an estuarine sea anemone found in salt marshes along both the Atlantic and Pacific coastlines of North America (Har 2009). They possess only two embryonic cell layers (i.e. diploblastic) and possess a single oral opening that is surrounded by 12 to 16 tentacles (Renfer *et al*. 2010). They have a clearly defined pharynx and mesenteries that run along the oral-aboral axis and a physa at the aboral end used for digging (Layden *et al*. 2010). *N. vectensis* has rapidly gained popularity as a model organism because it is easy to rear in the lab, simple to manipulate, and has a wide range of available genomic resources (Putnam *et al*. 2007; Reitzel, Ryan and Tarrant 2012).

*N. vectensis* microbiome 16S rRNA amplicon high-throughput sequencing has increased our understanding of the microbial community diversity which is crucial for understanding nutrient cycling, defense against diseases, thermoregulation, and the ability to rapidly adapt to environmental stressors (Rosenberg *et al*. 2007; Mortzfeld *et al*. 2016). Previous studies on wild and lab raised *N. vectensis* using 16S rRNA gene sequencing, gene expression analysis, and whole genome sequencing found that they primarily associate with the bacteria phylum : Bacteroides, Chloroflexi, Cyanobacteria, Firmicutes, Proteobacteria, and Spirochetes (Har *et al*. 2015a). Other studies have also found that the *N. vectensis* microbiome can be dynamic with diel lighting and developmental stage heavily influencing the microbial community composition (Mortzfeld *et al*. 2016; Domin et al. 2018; Leach, Carrier and Reitzel 2019). The current information on *N. vectensis* suggests that the microbiome is a highly dynamic and complex community that responds to environmental cues. However, these patterns were observed in the whole organism, which may obscure important niche specific bacteria compartments within *N. vectensis*.

In this study, we distinguished body region-specific bacterial associations present within *N. vectensis* by using 16S rRNA gene amplicon sequencing of the three body compartments: the capitulum (pharynx + tentacles), mesenteries, and physa (Fig. 1). We found that the capitulum possesses significantly different bacterial community abundances compared with the mesenteries and physa and that spirochetes are the dominant bacterial phylum present in the capitulum. Spirochetes were previously identified in *N. vectensis*; however, their compartment specific nature was unknown. The work presented better clarifies our understanding of bacteria community diversity in *N. vectensis* and sheds light on the importance of compartmental sampling in microbiome studies.

**Fig 1.**
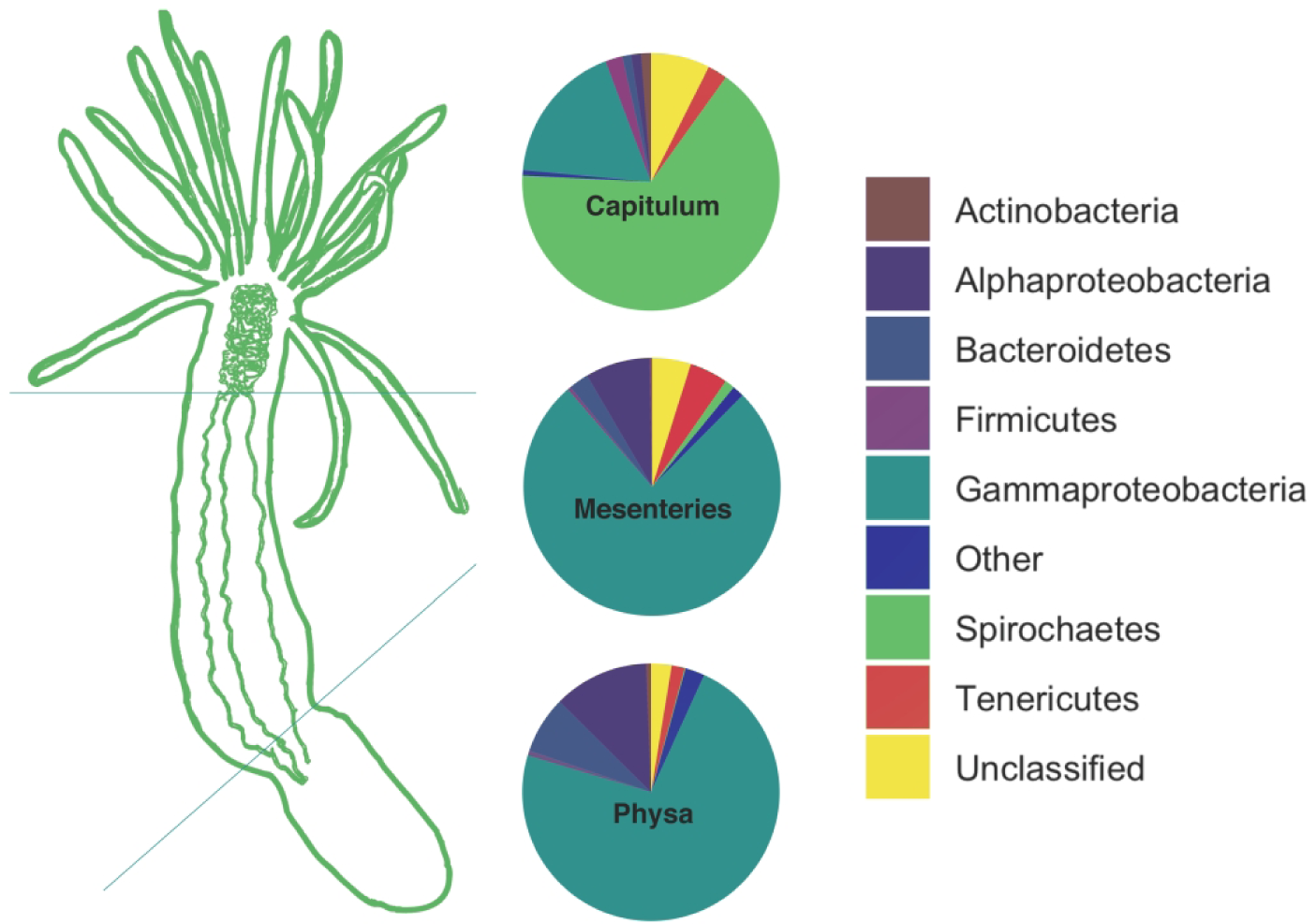
Compartmentalization of the microbiome in *N. vectensis*. The microbiome of *N. vectensis* was divided into three tissue compartments for this study: the capitulum, mesenteries, and physa. The capitulum had a bacterial community dominated by Spirochetes. The mesenteries and physa had a bacterial community dominated by Gammaproteobacteria and Alphaproteobacteria, with very little presence of Spirochetes. 338×218mm (300 × 300 DPI)

## Methods

### Nematostella vectensis Culture and Care

The individuals used for this study are lab cultured animals that have been housed at the Rosenstiel School of Marine and Atmospheric Science (RSMAS) Cnidarian Immunity Lab since 2016. All animals were kept in the dark within covered, glass dishes filled with non-circulating 11 parts per thousand (ppt), 2-micron filtered seawater (FSW) at room temperature (∼23.8°C). Two months prior to the beginning of this study, *N. vectensis* adults were placed into a single glass dish and fed mussels three days a week, and *Artemia* five days a week for two months in order to increase their size. Water changes were conducted an hour after each mussel feeding. One week before beginning the trisection (see section below), 12 of the largest individuals were transferred individually into a 12-well plate with sterilized FSW and starved for seven days to minimize microbial carryover from food. Previous studies have also used a starving period of two days to eliminate digested food debris (Har *et al*. 2015b; Leach, Carrier and Reitzel 2019)

#### Trisection

The 12 adult *N. vectensis* were anesthetized using 7% MgCl_2_ solution and then aseptically trisected. The trisection resulted in the isolation of three tissue compartments from each anemone: (1) capitulum (pharynx and tentacles), (2) mesenteries, and (3) physa (Fig. 1). The scalpel and tweezers were both flame and ethanol sterilized between tissue divisions. Each cut was made as cleanly as possible to limit any cross-contamination of tissue compartments. The capitulum was removed first, followed by the mesenteries, then the physa. The cut for the capitulum and mesentery boundary was established as just below the visible pharynx. Some physa tissue was found in the mesentery compartment as the mesenteries were not directly removed from the body cavity. However, the physa compartment did not contain any mesentery material. Each sample was then stored in a 2 mL centrifuge tube with RNALaterTM (Sigma Aldrich, St. Louis, MO) and placed in a -80°C freezer. In total there were 18 samples (n=6 capitulum, n=6 mesenteries, and n=6 physa) that were further processed.

### 16S rRNA PCR and Sequence Library Preparation

Total DNA was extracted from 18 samples using the Qiagen(tm) DNeasy Powersoil Kit. The concentration and quality of the DNA extracts were assessed using a Nanodrop(tm) spectrophotometer (Thermo Fisher Scientific Inc., Waltham, MA). DNA was then stored in a -80°C freezer until library preparation. High-throughput 16S rRNA amplicon sequencing methods were adapted from the protocol developed by the Earth Microbiome Project (Thompson *et al*. 2017). In short, the V4 region of the 16S rRNA gene (515F-806R primers) was amplified using polymerase chain reaction (PCR) on 18 tissue samples (Apprill *et al*. 2015). A negative non-template control was also amplified to account for extraction and PCR contamination. A total of 50 µL of PCR reaction mixtures were prepared according to the following recipe: 23 µL PCR-grade water, 20 µL PCR master mix (2x), 1 µL 806R primer (10 µM), 1 µL barcoded 515F primer (10µM), and 5 µL template DNA. PCR was performed with the following program: 1 × 3 min at 94°C, 35 × (45 s at 94°C, 60 s at 50°C, 90 s at 72°C), 1 × 10 min at 72°C. Amplified DNA was stored at -20°C for one week. The PCR product was run on a 1.5% agarose gel to check for correct amplification size. Following the gel, PCR products were cleaned using AMPure XP Beads(tm) (Beckman Coulter Inc., Brea, CA). DNA concentrations were quantified using the Qubit dsDNA high sensitivity (HS) kit(tm) (Thermo Fisher Scientific Inc., Waltham, MA) and samples were diluted to 4 nM using PCR-grade water. Then 5 µL of each sample was pooled into a single 1.5 mL tube. The pooled DNA library was sent for sequencing on an Illumina MiSeq(tm) PE300 sequencer at the Center for Genome Technology (CGT) at the University of Miami’s Miller School of Medicine for quality control and sequencing. Raw Illumina paired-end reads have been deposited in the NCBI Sequence Read Archive (SRA) database under BioProject PRJNA610635.

### Post-Sequencing Processing and Statistical Analysis

Post-sequencing processing was conducted using the Quantitative Insights in Microbial Ecology 2 (QIIME2) v2019.1 pipeline (Bolyen *et al*. 2019). Demultiplexed paired-end reads were merged using DADA2 v1.10.0 and processed to yield amplicon sequence variants (ASVs) (Callahan *et al*. 2016). Within the DADA2 pipeline, forward sequences were truncated at 200 bp and reverse sequences were truncated at 180 bp. Taxonomy was assigned using the SILVA 132 99% classifier (https://data.qiime2.org/2019.10/common/silva-132-99-nb-classifier.qza) (Quast *et al*. 2013). Mitochondria and chloroplast sequences were filtered from the feature table. The Phyloseq R package v1.28.0 was used for data handling (McMurdie and Holmes 2013). The R package decontam v1.4.0 was used to eliminate microbial contaminants by removing all sequences that were more prevalent in the negative control than in the real samples (Davis *et al*. 2018). Normalization across samples was conducted using a variance stabilizing transformation (VST) on the data using DESeq2 v1.24.0 (Love, Huber and Anders 2014). To account for abundance and evenness of microbial taxa within the VST dataset, the Shannon-Weiner Diversity Index and Simpson’s Diversity Index were calculated for each compartment region using Phyloseq. DESeq2 v1.24.0 and the QIIME2 add-on, Analysis of Composition of Microbiomes (ANCOM), were used for microbial differential abundance analysis against tissue-compartments and differences were considered significant if they had a p-value < 0.05. Random forest classifier modeling was performed using randomForest in order to determine the main bacterial taxa driving tissue compartment composition (Liaw and Wiener 2002). For beta-diversity measurements, both Bray-Curtis and Aitchison (Euclidean distances of center-log ratio transformed counts) distance matrices were assessed as both are commonly used in microbial community studies. For Bray-Curtis distances, the VST-transformed data was used. For Aitchison distances, the R package Microbiome v1.6.0 was used to perform a center-log ratio transformation on the raw count data (Lahti *et al*. 2019). Vegan v2.5.6 was used to calculate Bray-Curtis and Euclidean distances and the *Adonis* function was used to perform PERMANOVA analyses in order to statistically assess dissimilarity in microbial community composition between tissue compartments (Oksanen *et al*. 2013). Ampvis2 v2.5.7 was used to generate a heat-map of the most abundant bacterial families according to tissue compartment (Andersen *et al*. 2018). All associated code can be found on GitHub: https://github.com/Abonacolta/Nvec_Compartmentalization.

### Mapping N. vectensis ASVs to a phylogenetic tree of Spirochetes

The 16S rRNA gene sequences from the phylum Spirochaetae were retrieved from NCBI for representatives from every major spirochete lineage in FASTA format. Using BLASTn, closely clustering (97%) sequences to the two spirochete ASVs recovered in this study were added to the FASTA file along with a Planctomycete sequence (Accession: X56305) as an outgroup, giving a total of 64 sequences. These sequences were aligned using MAFFT v7.450 (auto-setting) with the L-INS-i strategy (Katoh and Standley 2013). The alignment was trimmed to remove columns with gaps in more than 70% of the sequences or with a similarity score below 0.001 using TrimAl v1.2 (Capella-Gutiérrez, Silla-Martínez and Gabaldón 2009). A maximum-likelihood tree was constructed with RAxML v7.3.0 using the GTRCAT substitution model and 1000 sub-samplings to generate a best-fit tree (Stamatakis 2014). The final tree was visualized and refined in FigTree v 1.4.4 (Drummond *et al*. 2012).

## Results and Discussion

### The body regions of N. vectensis exhibited distinct bacterial associations

We had an average of 113,758 reads per sample and there was a total of 620 ASVs with five ASVs removed as they were likely contaminants. The microbial composition of the capitulum (n= 6) is distinct from the mesenteries (n= 6) and physa (n= 6; Fig. 1). With the mesenteries showing a similar bacterial community composition as the physa. Overall the microbiomes were consistent between replicates, however, one individual harbored a different compartmental microbiome from the others in this study (Figs. 3, S1, S2, & S3). Spirochetes (predominantly from 1 ASV) from the family Spirochaetaceae made up ∼66% of the bacterial community within the capitulum of *N. vectensis*, but only 1.2% and 0.1% of the community in the mesenteries and physa, respectively (Fig. 2). This finding is similar to the microbiome of the Crown-of-thorns Starfish, *Acanthaster planci*, which found a high relative abundance (43-64%) of spirochetes (predominantly from 1 ASV) only within the body wall compartment. Prior to this Alphaproteobacteria was thought to be the most abundant bacterial phylum across the sea stars (Høj *et al*. 2018).

**Fig 3.**
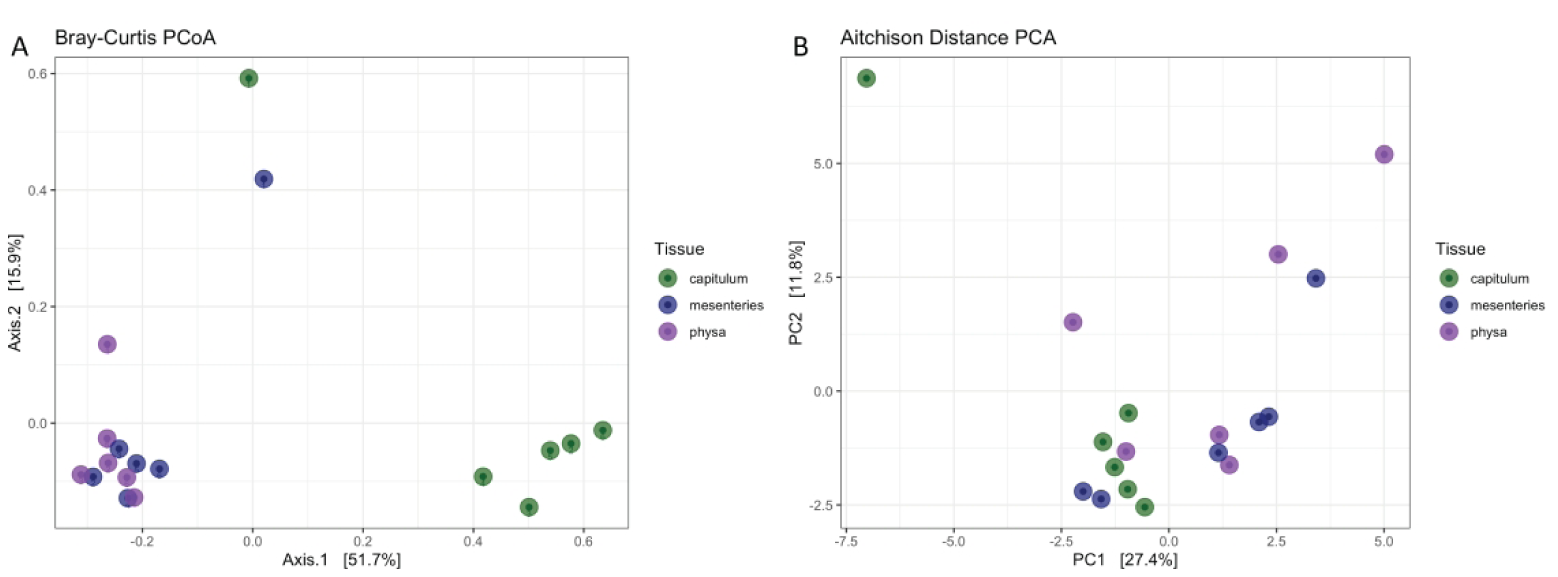
Beta diversity measures between tissue regions of *N. vectensis*. (A) Bray-Curtis PCoA of *N. vectensis* bacterial communities after a variance stabilizing transformation. Capitulum samples clustered tightly, while physa and mesentery communities clustered closely together. (B) Aitchison PCA (Euclidean distance of center-log ratio transformed sample counts) of *N. vectensis* bacterial communities. Again, most capitulum samples clustered closely together, however they were also clustered with mesenteries and physa samples. 355×127mm (300 × 300 DPI)

In the mesenteries a larger prevalence of Gammaproteobacteria was found (Fig. 1 & S2), which may play an important role in microbial community structuring in juvenile *N. vectensis* (Domin *et al*. 2018). The physa of *N. vectensis* was dominated by the classes Gammaproteobacteria, Alphaproteobacteria, and Bacteroidetes (Fig. S2). Similarly, previous studies on the microbiome of lab-raised *N. vectensis* found that Proteobacteria (including the classes Gammaproteobacteria and Alphaproteobacteria) was the most abundant bacterial phylum, with Firmicutes, Planctomycetes, Tenericutes, Verrucomicrobia, and Spirochetes having lower, but still significant, abundances (Har *et al*. 2015a; Mortzfeld *et al*. 2016; Leach, Carrier and Reitzel 2019). In addition to the physa, the mesenteries were also dominated by the class Gammaproteobacteria, among them Oceanospirillales and Pseudomonadales were the primary orders, with Pseudomonadales being more abundant than Oceanospirillales (Fig. S3). Among bacterial families, Pseudomonadaceae had the highest relative abundance in both the mesenteries and physa (∼36.5% and ∼38.1%, respectively) (Fig. 2). This result is in line with a fluorescent *in situ* hybridization study that revealed that Proteobacteria and Pseudomonas occur predominantly in aggregates along the walls of the mesenteries (Har 2009).

Interestingly, one study found that as *N. vectensis* develops, Betaproteobacteria, Actinobacteria, and Bacteroidetes decrease, while Spirochetes become one of the most abundant phyla in the adult organism (Mortzfeld *et al*. 2016). The development and growth of the capitulum during this time span may explain the high relative abundance of spirochetes that we saw in the capitulum. Spirochetes may play an active role in the capitulum development of *N. vectensis* and their ability to establish dominance over the Proteobacteria present throughout the rest of *N. vectensis* implies a competitive advantage on the part of the Spirochete. Of note is that Spirochetes are not found in high abundance in artemia or mussel, the two food sources for the *Nematostella* used in this study (Quiroz *et al*. 2015; Weingarten, Atkinson and Jackson 2019).

Spirochetes have been identified as symbionts in bivalves, corals, oligochaetes, and termites (as epibionts on intestinal protists) where they perform reductive acetogenesis and nitrogen fixation (Margulis, Nault and Sieburth 1991; Lilburn, Schmidt and Breznak 1999; Campbell and Cary 2001; Ohkuma *et al*. 2015). In cnidarians, spirochetes have been identified in a wide range of cnidarians including red coral, *Corallium rubrum, Hydra vulgaris* and the octocoral, *Lobophytum pauciflorum (*Fig. 4; Van De Water *et al*. 2016; Wessels *et al*. 2017; Hufnagel and Myhal 1977). In *L. pauciflorum* spirochetes are hypothesized to play a role in nitrogen fixation and microbial community (Van De Water *et al*. 2016; Wessels *et al*. 2017) however, in other cnidarians, their function is not well understood. The predominance of spirochetes as symbionts in multiple cnidarians suggests a potential conserved benefit to the cnidarian hosts. In the future, testing the role of spirochetes in nitrogen fixation in the capitulum of *N. vectensis* may inform our understanding of these potential symbionts.

**Fig 4.**
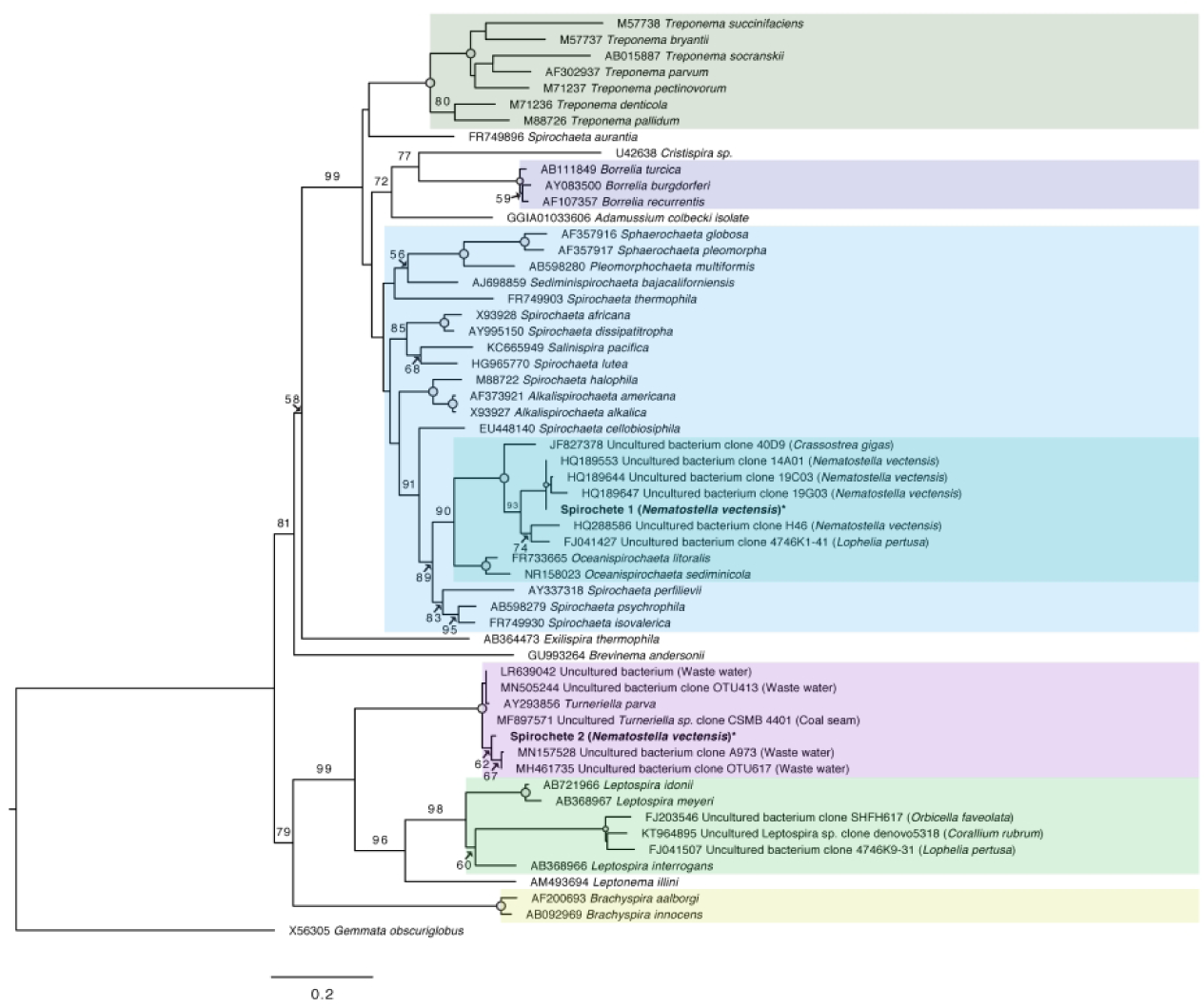
Best-fit maximum likelihood tree based on 16S rRNA sequences depicting the phylogenetic relationship of the spirochetes recovered from the capitulum of *Nematostella vectensis*. This tree was constructed using RAxML v7.3.0 with 1000 sub samplings. Bootstrap values (> 50%) are shown. Nodes with 100% bootstrap support are indicated by open circles. GenBank accession numbers are listed before each sequence’s name. The host/habitat of uncultured species is in parentheses. Spirochetes from this study are in bold. Shading corresponds to major clades within the phylum Spirochaetae. *Gemmata obscuriglobus* was used as an outgroup to root the tree. The predominant spirochete ASV from this study (Spirochete 1) forms a strongly supported (100% bootstrap) monophyletic clade with spirochete sequences recovered from wild *N. vectensis*. Spirochete 2 clusters within the genus *Turneriella*.

### Differential microbial community structure between body regions

The taxonomic make-up of the microbiome of *N. vectensis* varied according to body region sampled. Comparing the differential abundance of taxa across tissue compartments using a random forest classifier showed that the phyla Lentisphaerae, Proteobacteria, and Spirochetes were the strongest predictors of tissue compartments. According to differential abundance analysis, Spirochetes and Fusobacteria were significantly more abundant in the capitulum compared to the other compartments (DESeq2; p-adj=0.0000 and 0.0003, respectively).

Lentisphaerae, Proteobacteria, and Chloroflexi were more abundant in the mesenteries compared to the capitulum (DESeq2; p-adj= 0.0000, 0.0058, and 0.0009, respectively). According to differential abundance analysis using ANCOM, Spirochetes and Bacteroidetes were the most differentially abundant phyla among tissue compartments (ANCOM; W = 16 & 14, respectively). Spirochetes was the only taxon to be considered significantly differentially abundant or as a strong predictor of body region in *N. vectensis* by all programs, emphasizing the significance of this association (Fig. S4). The differences in dominant taxa between the capitulum and the other two tissue compartments in *N. vectensis* highlight the need to use compartmentalization to better understand the roles of the microbiome within the holobiont – be that in metabolism, defense, or nutrient cycling.

The body compartments of *N. vectensis* sampled in this study exhibited distinct microbial community structures. Alpha-diversity metrics showed a significant difference in the biodiversity and species richness present in different tissue compartments. Species richness among the capitulum and the mesenteries and physa were significantly different (ANOVA; p = 0.023, p = 0.033, respectively; Fig. S5). The Simpson’s Index between the capitulum and the mesenteries and physa was also significantly different (ANOVA; p = 0.039, p = 0.022, respectively; Fig. S5). However, the Shannon Diversity Index did not show a significant difference in alpha-diversity among body regions (Fig. S5). Overall, the capitulum of *N. vectensis* showed lower species richness compared to the mesenteries and physa, which had very similar species richness across individuals. A similar pattern was identified in the Simpson’s Index, with the capitulum showing a lower species diversity, but with a higher variation between individuals. The lower species richness in the capitulum was likely driven by the dominance of the one single Spirochete ASV. Both Bray-Curtis Dissimilarity and Aitchison distance were used to assess the differences in the microbial communities between tissue compartments (Fig. 3). A multivariate homogeneity of groups dispersions analysis confirmed that an ANOVA test could appropriately be performed on both datasets (Koleff, Gaston and Lennon 2003). Bacterial communities between tissue compartments showed significant dissimilarity (PERMANOVA; permutations = 9999, p = 0.0315) according to the Bray-Curtis Dissimilarity Index. However, the same PERMANOVA test on the Aitchison distances found insignificant dissimilarity between tissue compartments (PERMANOVA; permutations = 9999, p = 0.3294).

### Phylogenetic analysis of N. vectensis Spirochetes

The majority of spirochetes found in the capitulum was from a single ASV (Spirochete 1) within the order Spirochaetales. In order to better understand the phylogenetic placement of the spirochetes within *N. vectensis*, we conducted a phylogenetic analysis of this ASV along with the other spirochete ASV (Spirochete 2) recovered in this study. Strong bootstrap support (93% bootstrap) placed Spirochete 1 as closely related to four spirochete ASVs recovered from wild *N. vectensis* from Har *et al*. 2015 supporting this spirochete as a consistent associate of the anemone (Fig. 4). The potential role of the spirochetes within the capitulum of *N. vectensis* should be investigated further as this relationship appears to be conserved, regardless of environment. These *N. vectensis* ASVs also showed a strong relation to a spirochete recovered from the cold-water coral, *Lophelia pertusa*. Together, these sequences formed a strongly supported (90%) monophyletic clade with the two representative species of the recently described genus *Oceanispirocheata*. The genus *Oceanispirochaeta* consists solely of two representatives, *O. litoralis* and *O. sediminicola*, which are Gram-negative, moderately halophilic, and obligately anaerobic (Subhash and Lee 2017).

Spirochete 2, along with several closely clustering wastewater ASVs, showed strong relation (100% bootstrap) to *Turneriella parva*, which is a Gram-negative, obligately aerobic spirochete with morphological similarities to *Leptospira* (Stackebrandt *et al*. 2013). Spirochetes associated with a diverse group of corals, including *Corallium rubrum, Orbicella faveolata, and Lophelia pertusa*, did form a well-supported (100% bootstrap) monophyletic clade within the genus *Leptospira*, despite the broad geographical ranges of these coral species (*O. faveolata* is a Caribbean species, *C. rubrum* is a Mediterranean species, & *L. pertusa* is a deep sea species) indicating a potentially conserved association of this bacteria with Cnidarians.

## Conclusion

In this study we characterized the microbiome of three distinct body compartments within *N. vectensis*. A capitulum-specific dominance of spirochetes was uncovered highlighting the possible importance of this bacteria in *N. vectensis* physiology. Previous bulk-sampling techniques could not untangle compartment-specific bacterial associations, which should be taken into consideration in future cnidarian studies. Specifically, the role of spirochetes within the capitulum of *N. vectensis* should be further considered in subsequent studies. Overall, our findings further establish spirochetes as consistent bacteria associates within *N. vectensis* and suggests that the functional role of this bacteria in the capitulum may be important for the anemone.

## Acknowledgements

Authors would like to thank members of the Cnidarian Immunity Lab for comments and suggestions as this project was developing. AMB: was supported by the RSMAS Small Undergraduate Research Grant Experience (SURGE) Award. NTK was supported by startup funds from University of Miami, Rosenstiel School of Marine and Atmospheric Sciences and by NSF Award #: 1951826

## Supplemental Material

**Fig S1.**
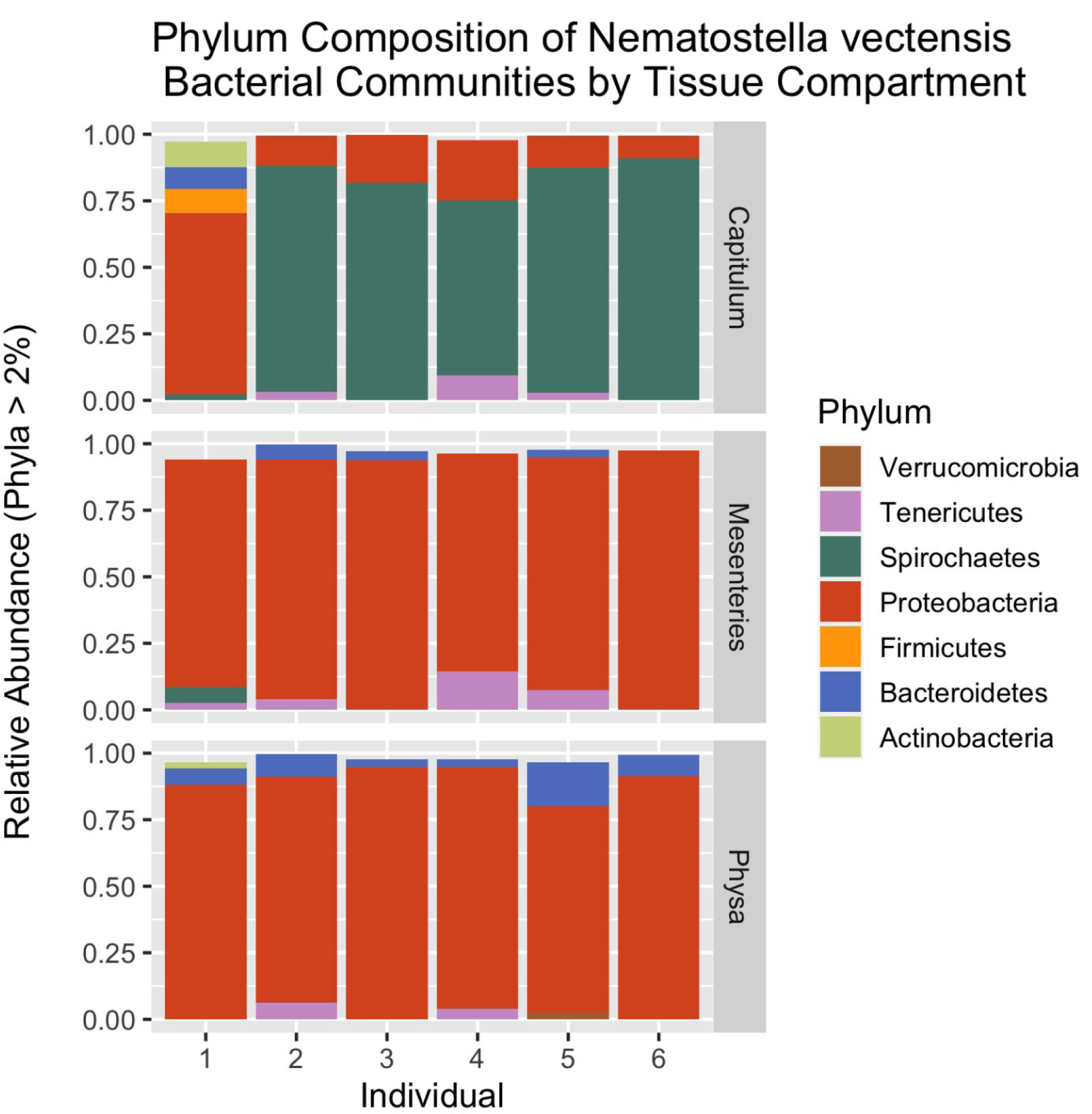
The relative abundance of bacterial phyla recovered from *N. vectensis* in individuals and tissue compartments. Only taxonomic phyla with a relative abundance above 2% were incorporated. Spirochetes dominated the bacterial communities in the majority of capitulum samples (Individual 1 again being an outlier). Proteobacteria dominated the bacterial communities of mesentery and physa samples. 529×529mm (72 × 72 DPI)

**Fig S2.**
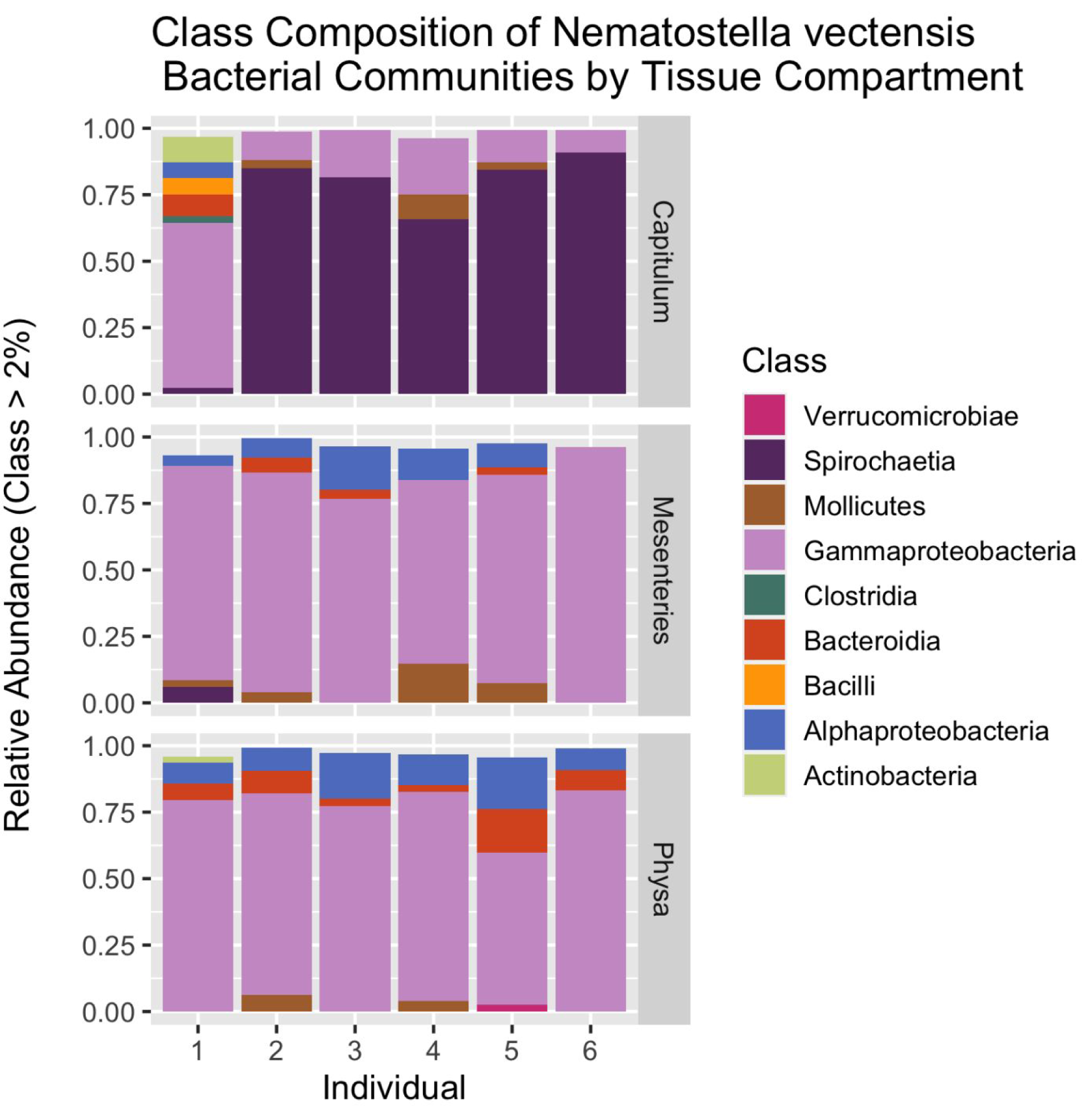
The relative abundance of bacterial classes recovered from *N. vectensis* in individuals and tissue compartments. Only taxonomic classes with a relative abundance above 2% were incorporated. Spirochaetia dominated the bacterial communities in the majority of capitulum samples (Individual 1 again being an outlier). Gammaproteobacteria dominated the bacterial communities of mesentery and physa samples.

**Fig S3.**
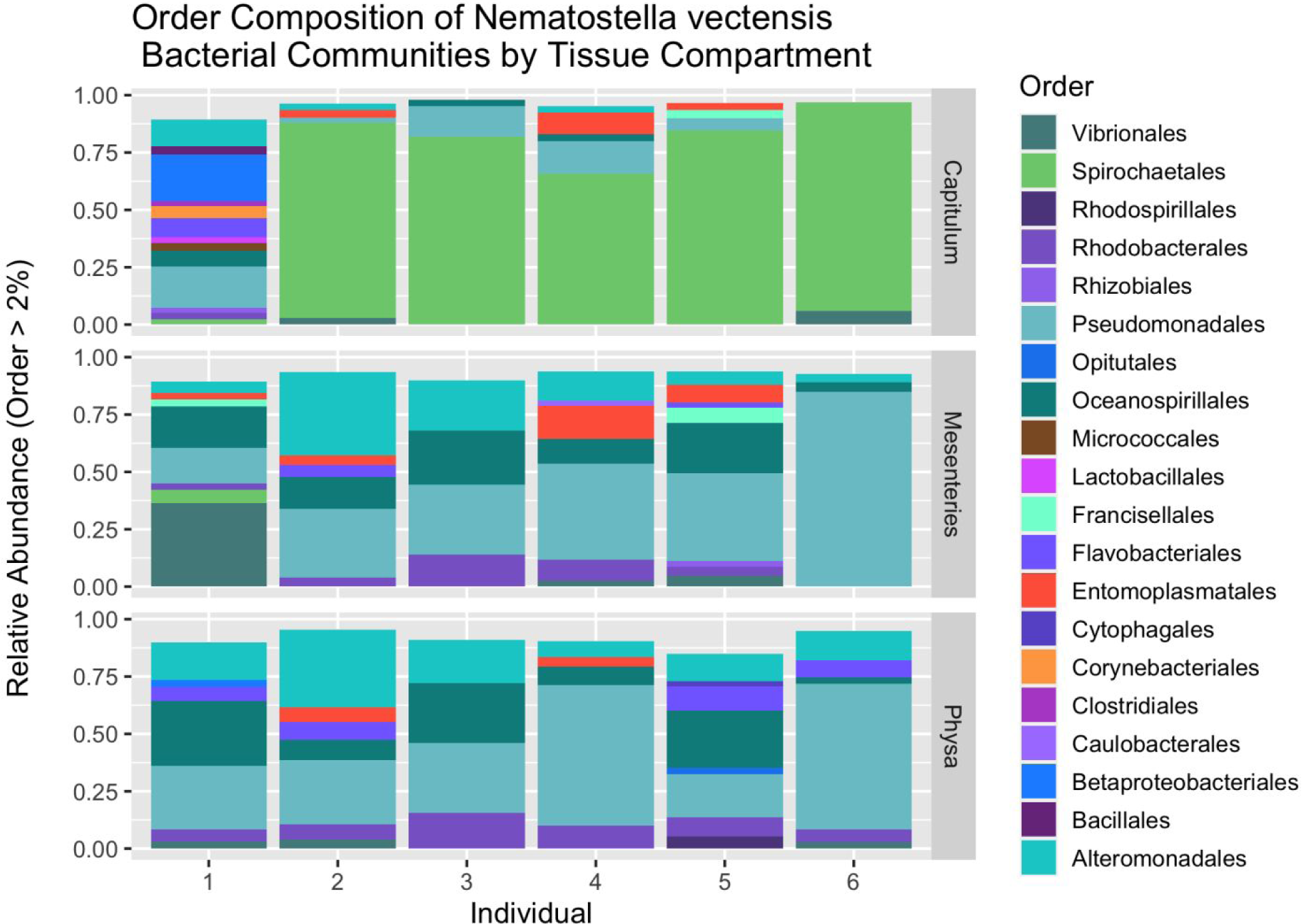
The relative abundance of bacterial orders recovered from *N. vectensis* in individuals and tissue compartments. Only taxonomic orders with a relative abundance above 2% were incorporated. The order Spirochaetales dominated the bacterial communities in the majority of capitulum samples (Individual 1 being an outlier). The mesentery and physa body regions showed more variety in their composition with the orders Pseudomonadales, Alteromonadales, and Oceanospirillales making up the majority of most samples. 740×529mm (72 × 72 DPI)

**Fig S4.**
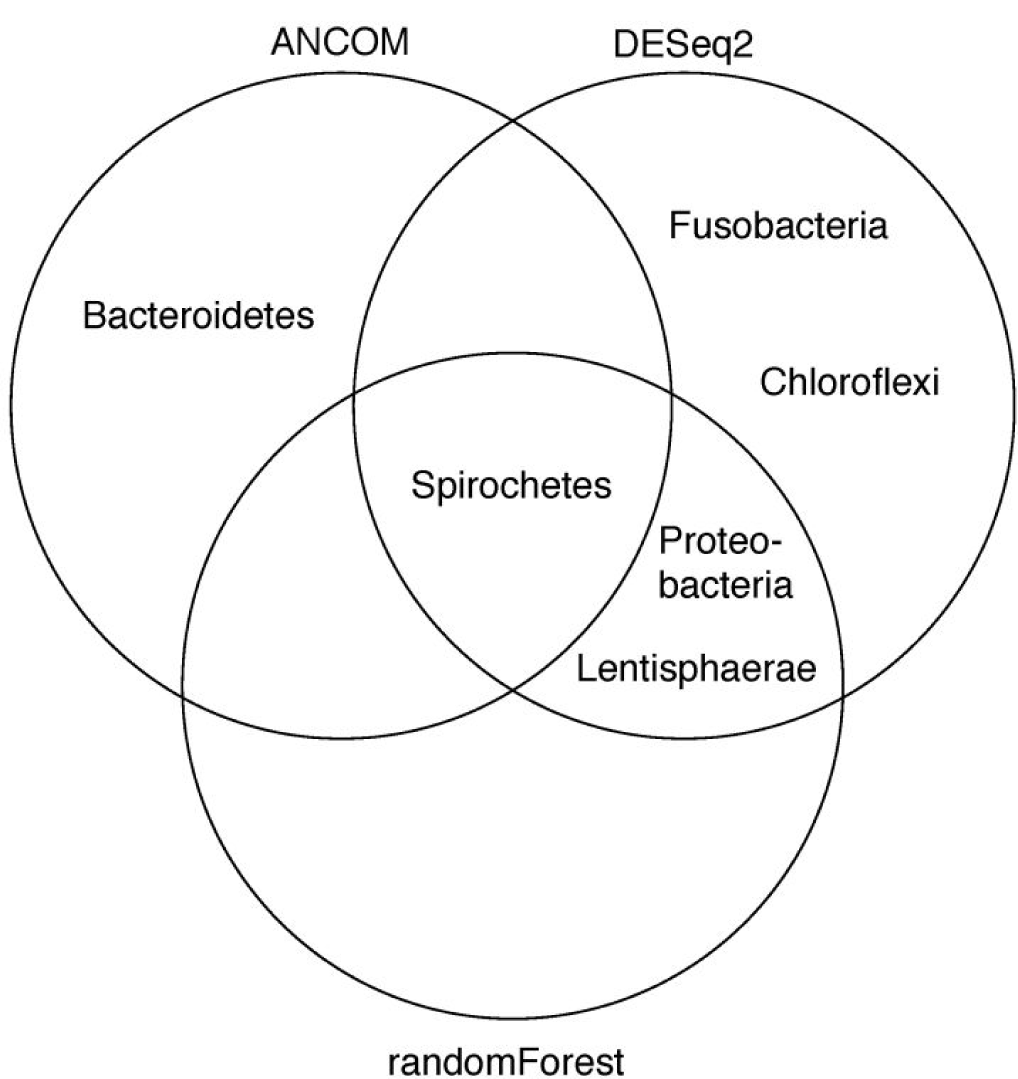
Venn diagram of significant taxa by program. Taxa with significant differential abundance between body regions were determined by ANCOM and DESeq2. Taxa which were shown to be the strongest predictor of body compartment by randomForest are also listed. Spirochetes was the only taxon to be considered significantly differentially abundant or as a strong predictor of body region in *N. vectensis* by all programs. 338×190mm (300 × 300 DPI)

**Fig S5.**
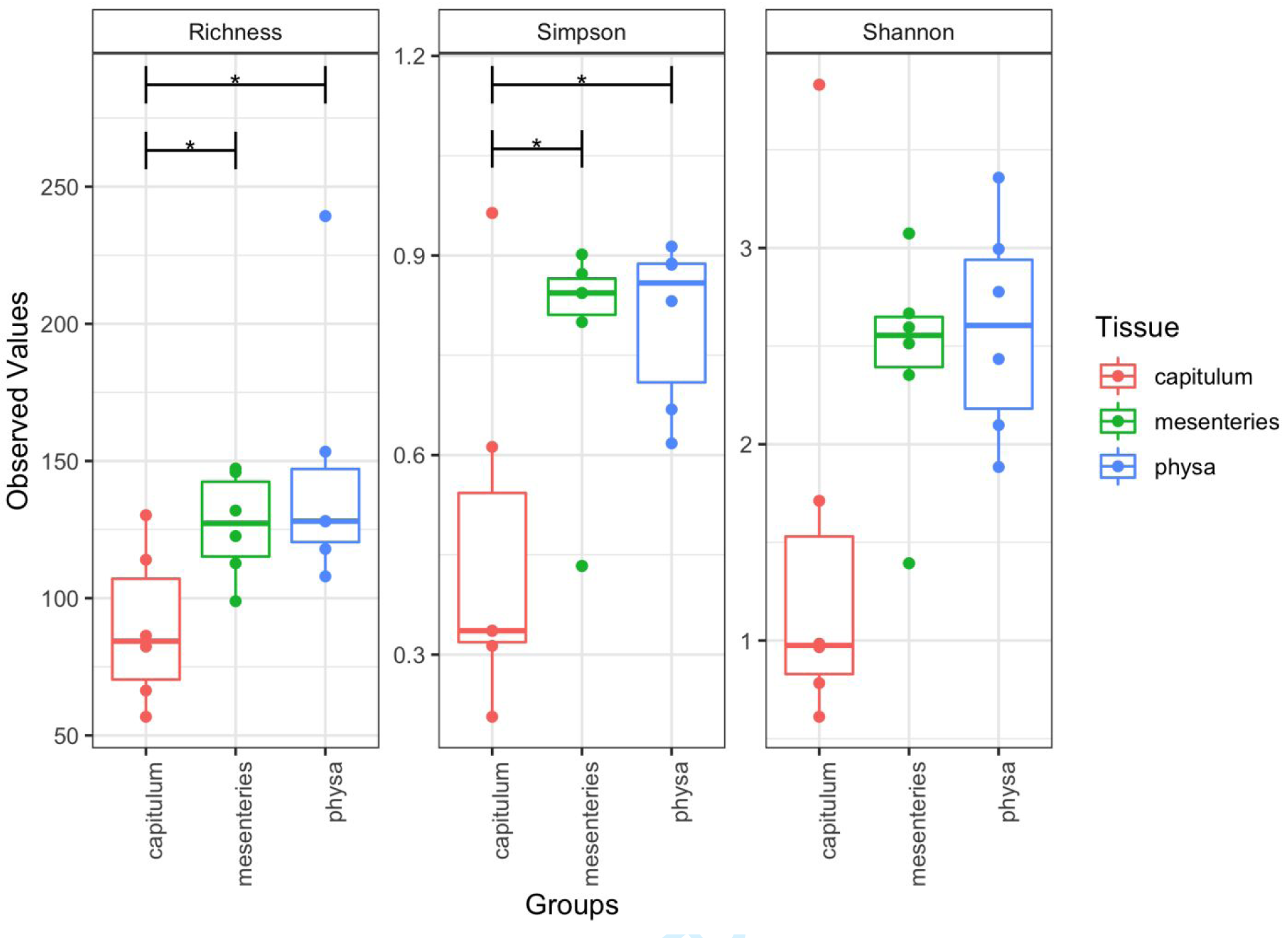
Bacterial community Alpha diversity measures between tissue regions. Species richness between the capitulum, mesenteries, and physa was significantly different (ANOVA; p=0.023, p=0.033, respectively). The Simpson Alpha Diversity Index between the capitulum, mesenteries and physa was significantly different (ANOVA; p=0.039, p=0.022, respectively). The Shannon Diversity Index did not show a significant difference in the bacterial communities between body regions.

## References

Aagaard K, Ma J, Antony KM et al. The Placenta Harbors a Unique Microbiome. Sci Transl Med 2014;6, DOI: 10.1126/scitranslmed.3009864.

Ainsworth TD, Thurber RV, Gates RD. The future of coral reefs: a microbial perspective. Trends Ecol Evol 2010;25:233–40.

Apprill A, Mcnally S, Parsons R et al. Minor revision to V4 region SSU rRNA 806R gene primer greatly increases detection of SAR11 bacterioplankton. Aquat Microb Ecol 2015;75:129–37.

Apprill A, Weber LG, Santoro AE. Distinguishing between Microbial Habitats Unravels Ecological Complexity in Coral Microbiomes. mSystems 2016;1:e00143–16.

Bolyen E, Rideout JR, Dillon MR et al. Reproducible, interactive, scalable and extensible microbiome data science using QIIME 2. Nat Biotechnol 2019;37:852–7.

Bourne DG, Morrow KM, Webster NS. Insights into the Coral Microbiome: Underpinning the Health and Resilience of Reef Ecosystems. Annu Rev Microbiol 2016;70:317–40.

Callahan BJ, Mcmurdie PJ, Rosen MJ et al. DADA2: High resolution sample inference from Illumina amplicon data. Nat Methods 2016;13:581–3.

Campbell BJ, Cary SC. Characterization of a Novel Spirochete Associated with the Hydrothermal Vent Polychaete Annelid, Alvinella pompejana. Appl Environ Microbiol 2001;67:110–7.

Capella-Gutiérrez S, Silla-Martínez JM, Gabaldón T. trimAl: A tool for automated alignment trimming in large-scale phylogenetic analyses. Bioinformatics 2009;25:1972–3.

Davis NM, Proctor DiM, Holmes SP et al. Simple statistical identification and removal of contaminant sequences in marker-gene and metagenomics data. Microbiome 2018;6:1–14.

Domin H, Zurita-Gutiérrez YH, Scotti M et al. Predicted bacterial interactions affect in vivo microbial colonization dynamics in Nematostella. Front Microbiol 2018;9:1–12.

Drummond AJ, Suchard MA, Xie D et al. Bayesian phylogenetics with BEAUti and the BEAST 1.7. Mol Biol Evol 2012;29:1969–73.

Foster KR, Schluter J, Coyte KZ et al. The evolution of the host microbiome as an ecosystem on a leash. Nature 2017;548:43–51.

Har JY. Introducing the starlet sea anemone Nematostella vectensis as a model for investigating. 2009.

Har JY, Helbig T, Lim JH et al. Microbial diversity and activity in the Nematostella vectensis holobiont: Insights from 16S rRNA gene sequencing, isolate genomes, and a pilot-scale survey of gene expression. Front Microbiol 2015a;6, DOI: 10.3389/fmicb.2015.00818.

Har JY, Helbig T, Lim JH et al. Microbial diversity and activity in the Nematostella vectensis holobiont: Insights from 16S rRNA gene sequencing, isolate genomes, and a pilot-scale survey of gene expression. Front Microbiol 2015b;6, DOI: 10.3389/fmicb.2015.00818.

Høj L, Levy N, Baillie B et al. Crown-of-Thorns Sea Star Acanthaster cf. solaris Has Tissue-Characteristic Microbiomes with Potential Roles in Health and Reproduction. Appl Environ Microbiol 2018;84:1–18.

Hooper L V., Littman DR, Macpherson AJ. Interactions between the microbiota and the immune system. Science (80-) 2012;336:1268–73.

Hufnagel LA, Myhal ML. Observations on a Spirochaete Symbiotic in Hydra. Trans Am Microsc Soc 1977;96:406–11.

Katoh K, Standley DM. MAFFT multiple sequence alignment software version 7: Improvements in performance and usability. Mol Biol Evol 2013;30:772–80.

Koleff P, Gaston KJ, Lennon JJ. Measuring beta diversity for presence-absence data. J Anim Ecol 2003;72:367–82.

Layden MJ, Meyer NP, Pang K et al. Expression and phylogenetic analysis of the zic gene family in the evolution and development of metazoans. Evodevo 2010;1:1–16.

Leach WB, Carrier TJ, Reitzel AM. Diel patterning in the bacterial community associated with the sea anemone Nematostella vectensis. Ecol Evol 2019;9:9935–47.

Liaw A, Wiener M. Classification and Regression by randomForest. R News 2002;2:18–22.

Lilburn TG, Schmidt TM, Breznak JA. Phylogenetic diversity of termite gut spirochaetes. Environ Microbiol 1999;1:331–45.

Love MI, Huber W, Anders S. Moderated estimation of fold change and dispersion for RNA-seq data with DESeq2. Genome Biol 2014;15:1–21.

Margulis L, Nault L, Sieburth JM. Cristispira from oyster styles: complex morphology of large symbiotic spirochetes. Symbiosis 1991;11:1–17.

McMurdie PJ, Holmes S. Phyloseq: An R Package for Reproducible Interactive Analysis and Graphics of Microbiome Census Data. PLoS One 2013;8, DOI: 10.1371/journal.pone.0061217.

Mortzfeld BM, Urbanski S, Reitzel AM et al. Response of bacterial colonization in Nematostella vectensis to development, environment and biogeography. Environ Microbiol 2016;18:1764–81.

Ohkuma M, Noda S, Hattori S et al. Acetogenesis from H2 plus CO2and nitrogen fixation by an endosymbiotic spirochete of a termite-gut cellulolytic protist. Proc Natl Acad Sci U S A 2015;112:10224–30.

Putnam NH, Srivastava M, Hellsten U et al. Sea anemone genome reveals ancestral eumetazoan gene repertoire and genomic organization. Science (80-) 2007;317:86–94.

Quast C, Pruesse E, Yilmaz P et al. The SILVA ribosomal RNA gene database project: Improved data processing and web-based tools. Nucleic Acids Res 2013;41:590–6.

Quiroz M, Triadó-Margarit X, Casamayor EO et al. Comparison of Artemia–bacteria associations in brines, laboratory cultures and the gut environment: a study based on Chilean hypersaline environments. Extremophiles 2015;19:135–47.

Rädecker N, Pogoreutz C, Voolstra CR et al. Nitrogen cycling in corals: The key to understanding holobiont functioning? Trends Microbiol 2015;23:490–7.

Reitzel AM, Ryan JF, Tarrant AM. Establishing a model organism: A report from the first annual Nematostella meeting. BioEssays 2012;34:158–61.

Renfer E, Amon-Hassenzahl A, Steinmetz PRH et al. A muscle-specific transgenic reporter line of the sea anemone, Nematostella vectensis. Proc Natl Acad Sci U S A 2010;107:104–8.

Ricci F, Rossetto Marcelino V, Blackall LL et al. Beneath the surface: Community assembly and functions of the coral skeleton microbiome. Microbiome 2019;7:1–10.

Robbins SJ, Singleton CM, Chan CX et al. A genomic view of the reef-building coral Porites lutea and its microbial symbionts. Nat Microbiol 2019:1–11.

Rosenberg E, Koren O, Reshef L et al. The role of microorganisms in coral health, disease and evolution. Nat Rev Microbiol 2007;5:355–62.

Rossbach S. Tissue-Specific Microbiomes of the Red Sea Giant Clam Tridacna maxima Highlight Differential Abundance of Endozoicomonadaceae. 2019;10, DOI: 10.3389/fmicb.2019.02661.

Stackebrandt E, Chertkov O, Lapidus A et al. Genome sequence of the free-living aerobic spirochete Turneriella parva type strain (HT), and emendation of the species Turneriella parva. Stand Genomic Sci 2013;8:228–38.

Stamatakis A. RAxML version 8: A tool for phylogenetic analysis and post-analysis of large phylogenies. Bioinformatics 2014;30:1312–3.

Subhash Y, Lee SS. Description of oceanispirochaeta sediminicola gen. Nov., sp. nov., an obligately anaerobic bacterium isolated from coastal marine sediments, and reclassification of spirochaeta litoralis as oceanispirochaeta litoralis comb. nov. Int J Syst Evol Microbiol 2017;67:3403–9.

Sweet MJ, Croquer A, Bythell JC. Bacterial assemblages differ between compartments within the coral holobiont. Coral Reefs 2011;30:39–52.

Thompson LR, Sanders JG, McDonald D et al. A communal catalogue reveals Earth’s multiscale microbial diversity. Nature 2017;551:457–63.

Vaishnava S, Behrendt CL, Ismail AS et al. Paneth cells directly sense gut commensals and maintain homeostasis at the intestinal host-microbial interface. Proc Natl Acad Sci U S A 2008;105:20858–63.

Van De Water JAJM, Melkonian R, Junca H et al. Spirochaetes dominate the microbial community associated with the red coral Corallium rubrum on a broad geographic scale. Sci Rep 2016;6:1–7.

Weingarten EA, Atkinson CL, Jackson CR. The gut microbiome of freshwater Unionidae mussels is determined by host species and is selectively retained from filtered seston. PloS One 2019;14:1–17.

Wessels W, Sprungala S, Watson SA et al. The microbiome of the octocoral Lobophytum pauciflorum: Minor differences between sexes and resilience to short-term stress. FEMS Microbiol Ecol 2017;93:1–13.

Wier AM, Sacchi L, Dolan MF et al. Spirochete Attachment Ultrastructure: Implications for the Origin and Evolution of Cilia. Biol Bull 2010;218:25–35.

